# Intolerance of uncertainty is associated with heightened responding in the prefrontal cortex during instructed outcome uncertainty of threat

**DOI:** 10.1101/2020.05.23.112268

**Authors:** Jayne Morriss, Tiffany Bell, Nicolò Biagi, Tom Johnstone, Carien M. van Reekum

## Abstract

Heightened responding to uncertain threat is considered a hallmark of anxiety disorder pathology. Here, we sought to determine if individual differences in self-reported intolerance of uncertainty (IU), a key transdiagnostic dimension in anxiety-related pathology, underlies differential recruitment of neural circuitry during instructed outcome uncertainty of threat (*n* = 42). In an instructed threat of shock task, cues signalled uncertain threat of shock (50%) or certain safety from shock outcomes. Ratings of arousal and valence, skin conductance response (SCR) and functional magnetic resonance imaging were acquired. Overall, participants displayed greater ratings of arousal and negative valence, SCR, and amygdala activation to uncertain threat vs. safe cues. IU was not associated with greater arousal ratings, SCR or amygdala activation to uncertain threat vs. safe cues. However, we found that high IU was associated with greater ratings of negative valence and greater activity in the medial prefrontal cortex and dorsomedial rostral prefrontal cortex to uncertain threat vs safe cues. These findings suggest that during instructed outcome uncertainty of threat, individuals high in IU rate uncertain threat as aversive and engage prefrontal cortical regions known to be involved in safety-signalling and conscious threat appraisal. Taken together, these findings highlight the potential of IU in modulating safety-signalling and conscious appraisal mechanisms in situations with instructed outcome uncertainty of threat, which may be relevant to models of anxiety-related pathology.

## Introduction

In everyday life, we often experience uncertainty, and will typically try to minimise or resolve it, in order to reduce anxiety and stress (Grupe & Nitschke, 2013; Morriss, Gell, & van Reekum, 2019; Peters, McEwen, & Friston, 2017). Individuals who score high in self-reported intolerance of uncertainty (IU) tend to find uncertainty particularly distressing (Carleton, 2016a, 2016b; Dugas, Buhr, & Ladouceur, 2004; Freeston, Rhéaume, Letarte, Dugas, & Ladouceur, 1994). IU is considered a transdiagnostic dimension, as high levels of IU are observed across many mental health disorders with an anxiety component such as anxiety, depression, post-traumatic stress and obsessive-compulsive disorder (Gentes & Ruscio, 2011; Mahoney & McEvoy, 2012). On this basis, there has been a surge in IU-related research in the field of anxiety over the last decade (McEvoy, Hyett, Shihata, Price, & Strachan, 2019; Tanovic, Gee, & Joormann, 2018).

Despite progress in understanding the aetiology of IU, there still remain gaps in the literature as to how IU modulates neural circuitry associated with the processing of outcome uncertainty of threat (i.e, whether a threat such as an aversive stimulus will occur or not) (Tanovic et al., 2018). Only a handful of studies to date have examined how IU is correlated with neural circuitry during the anticipation of outcome uncertainty of threat (Morriss, Christakou, & Van Reekum, 2015; Schienle, Köchel, Ebner, Reishofer, & Schäfer, 2010; Simmons, Matthews, Paulus, & Stein, 2008; Somerville et al., 2013). In tasks where participants have been instructed about the likelihoods of negative or neutral pictures (i.e., visual cues or countdowns signal the occurrence of a negative or neural picture), individuals high in IU, relative to low IU, have been shown to exhibit heightened amygdala and insula activity to cues signalling unpredictable negative pictures (Schienle et al., 2010; Shankman et al., 2014), and exaggerated amygdala activity to negative pictures following unpredictable countdowns (Somerville et al., 2013). Whilst previous work has provided a starting point for understanding how IU modulates neural circuitry to outcome uncertainty of threat, further research is needed to assess the robustness and generalisability of IU-related effects. For example, previous fMRI research on the relationship between IU and instructed outcome uncertainty of threat has primarily used negative and neutral picture stimuli with a wide range of content. It is important to establish whether similar patterns of neural activation to instructed outcome uncertainty of threat would be observed for individuals with high IU in response to other stimuli commonly used to evoke anxious states such as mild electric shock. Understanding how IU modulates neural circuitry in relation to instructed outcome uncertainty of threat will further clarify the role of IU in neurobiological (Grupe & Nitschke, 2013; Peters et al., 2017) and clinical models of anticipation and uncertainty in anxiety (Carleton, 2016b; Shihata, McEvoy, Mullan, & Carleton, 2016).

To assess the relationship between self-reported IU and anticipatory responding during instructed outcome uncertainty of threat, we measured event-related functional magnetic resonance imaging (fMRI), skin conductance response (SCR) and, arousal and valence ratings while participants performed an instructed threat of shock task. To induce a sense of uncertain threat, a cue signalled whether a mild electric shock to the finger would occur (50% of the time) (i.e. participants were told that they would sometimes receive a shock at the end of the cued trial). Trials were 9 seconds in length (1 second cue, 8 second anticipatory period), to allow us to examine phasic and sustained threat/safety related activity.

We hypothesized that, during the instructed threat of shock task, we would observe typical patterns of phasic and sustained activation in circuitry associated with the processing of threat and safety (Etkin, Egner, & Kalisch, 2011; Levy & Schiller, 2021; Mechias, Etkin, & Kalisch, 2010; Morriss et al., 2018; Tashjian, Zbozinek, & Mobbs, 2021), i.e. (1) greater activation in the amygdala, putamen, caudate, insula and rostral prefrontal cortex to threat trials and (2) greater medial prefrontal cortex activity to safe trials. Moreover, we hypothesized that participants would display greater SCR to the threat vs. safe trials, and rate the threat trials as more negative and arousing than the safe trials.

Based on past research (Morriss et al., 2015; Schienle et al., 2010; Shankman et al., 2014; Simmons et al., 2008; Somerville et al., 2013), we hypothesised that higher IU would be associated with: (1) greater phasic and sustained activation in the amygdala and insula during threat, relative to safe trials, and (2) modulation of the medial prefrontal cortex during threat, relative to safe trials. Given the shortage of research on the relationship between IU and activation in the medial prefrontal cortex, we did not hypothesise a particular direction of effect. Lastly, we hypothesised that higher IU would be associated with greater SCR to the threat vs. safe trials, as well as higher ratings of negativity and arousal to the threat vs. safe trials.

We tested the specificity of the involvement of IU by comparing it with broader measures of anxiety (for discussion see (Morriss, Christakou, & Van Reekum, 2016), in this case the Spielberger State-Trait Anxiety Inventory, Trait Version (STAI-T) (Spielberger, Gorsuch, Lushene, Vagg, & Jacobs, 1983).

## Methods

### Participants

Forty-two female volunteers were recruited from the local area through advertisements (*M* age= 33 yrs, *SD* age= 7.33 yrs; 31 White Northern European, 6 White Non-Specified Region, 3 White Southern European, 1 Multi-ethnic, 1 White Eastern European). All participants reported being right-handed, having normal or corrected to normal vision, being medication-free and having no prior history of brain injury. We did not collect information from participants regarding current or previous history of mental health diagnoses. We selected female participants because the study was part of a larger programme of research examining the role of conspecifics (i.e. romantic partner, friend) in the processing of threat (Morriss, Bell, Johnstone, van Reekum, & Hill, 2019).

Participants provided written informed consent and received a picture of their brain and £15 for their participation. The University of Reading’s Research Ethics Committee approved the study protocol.

### Instructed threat of shock task

Participants were required to passively view cues that represented either threat of shock or safety from shock. Only two cues were presented, an uncertain threat cue where there was 50% chance of receiving a shock and a safety cue where there was 0% chance of receiving a shock. Each trial consisted of: a white cue (e.g. X, O, D, T) presented on a black background (1 second), a white fixation anticipation cue presented on a black background (8 seconds), a small circle cue signalling the end of the trial (1 second) and a black blank screen (4-6 seconds) (see Figure 1). In uncertain threat condition, the shock was administered with the end of trial cue 50% of the time. Participants completed 1 run of 36 trials (18 Threat, 18 Safe). To rule out any cue-specific effects, half the participants received X and O cues, whilst the other half received D and T cues.

**Figure 1:**
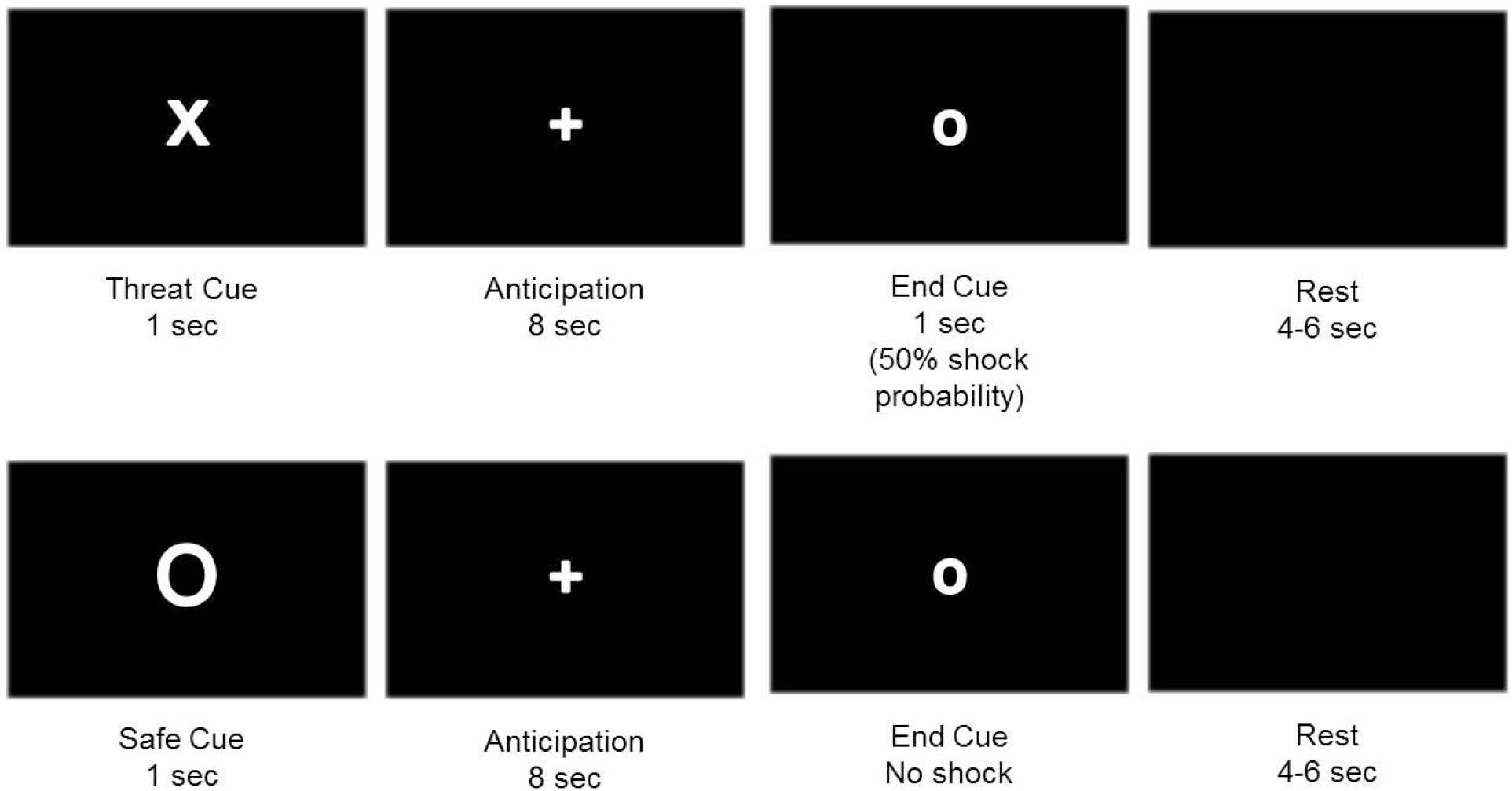
Image depicting instructed threat of shock task. Examples of threat (top row) and safe (bottom row) trial types. Participants were instructed on threat and safe contingencies before the start of the task.

At the beginning of the experiment four practise trials (2 Threat, 2 Safe) were presented without shock on a computer screen. While the participants were viewing these practise trials, the experimenter explained that one of the cues would be associated with shock and another would not. Additionally, the experimenter explained that there would be a waiting period and that if the shock was to occur, it would happen at the same time as the small circle cue (see Figure 1). The practise trials were given to allow the participants to experience the trial structure and timings. Before participants were placed in the scanner, they were instructed via a computer screen as to which cue would signal the possibility of a shock and which cue would signal no shock.

### Electric Shock

The possibility of receiving an unpleasant electrical shock to the index and middle finger of the right hand was used to induce threat. Electric shocks were delivered via an ADInstruments ML856 PowerLab 26T Isolated Stimulator using an MLADDF30 stimulating bar electrode with 30 mm spacing of 9 mm contacts. Each participant’s stimulation level was set by first exposing them to an electric stimulation of 1 mA (10 pulses at 50 Hz, with a pulse duration of 200 μs) and increasing the current in steps of 0.5 mA, up to a maximum of 10 mA. This continued until a suitable participant-specific threshold was found that was uncomfortable but not painful. This level was then used throughout the threat of shock task for that subject (electric stimulation level: M= 2.21 mA; SD= 1 mA).

### Procedure

Participants arrived at the laboratory and were informed of the experimental procedures. First, participants completed a consent form as an agreement to take part in the study. Second, participants completed questionnaires by pen and paper. Thirdly, participants were taken to the MRI unit, where they were presented with example trials from the instructed threat of shock task and the shock procedure was carried out. Next, we presented instructions about the task contingencies and then participants completed an instructed threat of shock task in the scanner whilst we concurrently recorded skin conductance. After scanning, participants rated the threat and safe cues from the instructed threat of shock task. Ratings were collected at the end of the scanning session rather than throughout the task because participants had electrodes attached to both hands.

### Questionnaires

To assess anxious disposition, we used Intolerance of Uncertainty (IU) (Freeston et al., 1994) and the Spielberger State-Trait Anxiety Inventory, Trait Version (STAI-T) (Spielberger et al., 1983). The IU scale comprises of 27 items on a 5-point Likert scale (possible range: 20-135) and captures aversion to uncertainty. The STAI-T comprises of 20 items on a 4-point Likert scale (possible range: 20-80) and captures trait anxiety. Both anxiety measures were normally distributed, showed good internal reliability and displayed similar ranges of scores to that of community samples (IU: *M* = 66.07; *SD* = 17.03; range = 34-102; α = .93; STAI-T; *M* = 40.92; *SD* = 10.31; range = 25-61; α = .91) (Julian, 2011; Khawaja & Yu, 2010). Importantly, at least half of the sample met STAI-T (>40) and IU (>60) scores that were comparable with clinical samples (Julian, 2011; Khawaja & Yu, 2010).

### Ratings

Participants rated the valence and arousal of the threat and safe cues using 9 point Likert scales ranging from 1 (Valence: negative; Arousal: calm) to 9 (Valence: positive; Arousal: excited). 1 participant did not complete the ratings, leaving 41 participants with ratings data.

### Skin conductance acquisition and reduction

Identical to previous work (Morriss et al., 2015), electrodermal recordings were obtained using AD Instruments (AD Instruments Ltd, Chalgrove, Oxfordshire) hardware and software. An ML138 Bio Amp connected to an ML870 PowerLab Unit Model 8/30 amplified the electrodermal signal, which was digitized through a 16-bit A/D converter at 1000 Hz. Electrodermal activity was measured during the scanning session with MRI-safe MLT117F Ag/AgCl bipolar finger electrodes filled with NaCl electrolyte paste (Mansfield R & D, St Albans, Vermont, USA) that were attached to the distal phalanges of the index and middle fingers of the left hand. A constant voltage of 22mVrms at 75 Hz was passed through the electrodes, which were connected to a ML116 GSR Amp.

Skin conductance responses (SCR) were scored when there was an increase of skin conductance level exceeding 0.03 microSiemens. The amplitude of each SCR was scored as the difference between the baseline (1 second average pre cue onset) and the maximum deflection (0.5-9 second post cue onset). Trials with no discernible SCRs were scored as zero. SCR magnitudes were calculated by averaging SCR values for each condition (Threat, Safe). Due to recording errors, 2 participants did not have SCR data, leaving 40 participants with SCR data.

### Ratings and SCR analysis

We conducted a 2 Condition (Threat, Safe) x IU ANCOVA for arousal ratings, valence ratings and SCR to the cues, where IU was entered as a continuous predictor variable. Any interaction with IU was followed up with pairwise comparisons of the means between the conditions for IU estimated at the specific values of + or - 1 SD of mean IU. This type of analysis with IU has been previously published elsewhere (Morriss et al., 2015, 2016).

We performed hierarchical regression analyses on the rating/SCR difference score measures that showed an effect with IU. This analysis served to assess IU-specific effects over and above shared variance with trait anxiety (STAI-T). We entered STAI-T in the first step and then IU in the second step.

### MRI

Participants were scanned with a 3T Siemens Trio using a 12 channel head coil (Siemens Inc., Erlangen, Germany). T2*-weighted gradient-echo, echo planar imaging (EPI) functional scans were acquired for the threat of shock task consisting of 281 volumes (TR = 2000 ms, TE = 30 ms, flip angle = 90°, FOV = 192 × 192 mm, 3 × 3 mm voxels, slice thickness 3 mm with an interslice gap of 1 mm, 30 axial slices, interleaved acquisition).

Following completion of the functional scan, structural and fieldmap scans were acquired, which comprised of a high-resolution T1-weighted anatomical scan (MP-RAGE, TR = 2020 ms, TE = 2.52 ms, flip angle = 90°, FOV = 256 × 256 mm, 1 x 1 x 1 mm voxels, slice thickness 1 mm, sagittal slices) and fieldmap (TR = 488 ms, TE 1 = 4.98 ms, TE 2 = 7.38 ms, flip angle = 60°, FOV = 256 × 256 mm, slice thickness 4 mm with an interslice gap of 4 mm, 30 axial slices).

### fMRI analysis

FMRI analyses were carried out in Feat version 5.98 as part of FSL (FMRIB’s Software Library, www.fmrib.ox.ac.uk/fsl). Brains were extracted from their respective T1 images by using the FSL Brain Extraction Tool (BET) (Smith, 2002). Distortion, slice timing and motion correction were applied to all extracted EPI volumes using FUGUE and MCFLIRT tools. Gaussian smoothing (FWHM 5mm) and a 50 second high pass temporal filter were applied.

A first-level GLM analysis was carried out for each functional scan. Separate regressors were specified for the experimental conditions of primary interest (Threat/Safety Cues) by convolving a binary boxcar function with an ideal haemodynamic response (HR), which corresponded to the length of each cue (1 second) or the entire trial period (9 seconds). Regressors for the end of trial period with and without shock and six motion parameters (3 rotation and 3 translation) were included to model out brain activity that was unrelated to the conditions of interest.

In two separate general linear models, we defined two main effect contrasts to reveal phasic and sustained threat/safety related activity: (1) Threat vs. Safety across the 1 second cue period, and (2) Threat vs. Safety across the whole 9 second trial period. All contrasts were normalized and registered to MNI standard space using FLIRT (Jenkinson, Bannister, Brady, & Smith, 2002). Second-level GLM analysis consisted of regressors for the group mean and demeaned IU scores using FSL’s OLS procedure. Whole-brain analysis was carried out with parametric statistics using cluster thresholding with a *z* = 2.3 and a corrected *p* < 0.05 (for fMRI results from non-parametric permutation tests, see supplementary material).

We performed hierarchical regression analyses on the resulting significant clusters that showed an association with IU to test for specificity of IU over STAI-T. We extracted % BOLD signal change difference scores from the relevant clusters and correlated these with the anxiety measures to test for IU-specific effects, by using STAI-T in the first step and then STAI-T and IU in the second step of hierarchical regression models.

## Results

### Ratings

Participants rated the threat cues as more negative (*M* = 4.78, *SD* = 1.77) and more arousing (*M* = 5.78, *SD* = 1.68) than the safe cue (*M* = 6.78, *SD* = 1.56 for valence and *M* = 2.90, *SD* = 2.11 for arousal respectively) [Condition (Valence): *F*(1,39) = 29.127, *p* < .001; Condition (Arousal): *F*(1,39) = 47.095, *p* < .001; see Figure 2A & B]. Higher IU was associated with significantly more negative ratings of the threat cues compared to the safe cues, *p* < .001 [Condition (Valence) x IU interaction: *F*(1,39) = 5.764, *p* = .021; see Figure 2C]. IU was not significantly related to the arousal ratings of the threat and safe cues [Condition (Arousal) x IU interaction: *F*(1,39) = .866, *p* = .358; see Figure 2D].

**Figure 2:**
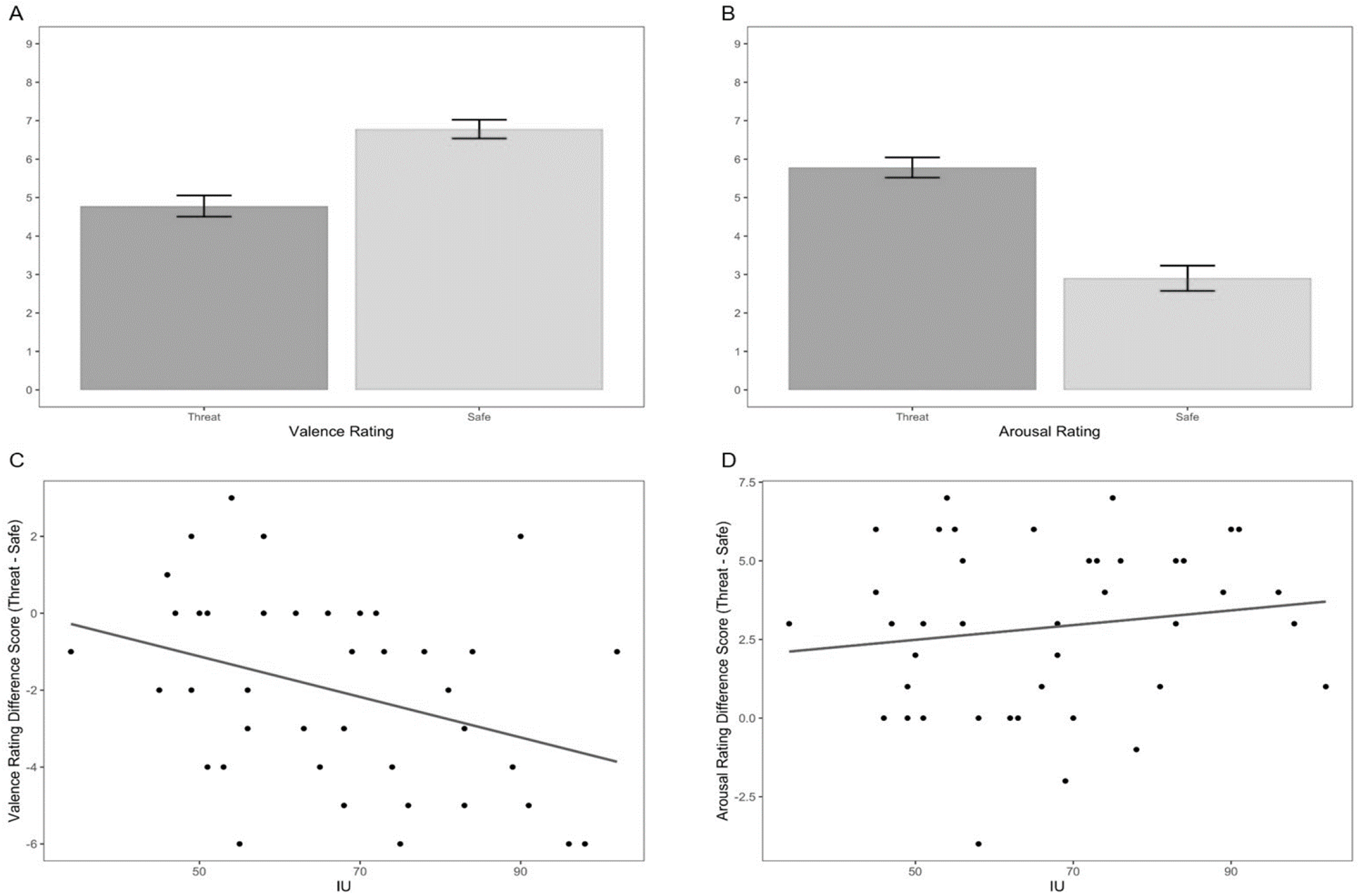
Bar graphs depicting valence and arousal ratings for threat and safe stimuli (A & B). Higher IU was significantly associated with rating the threat stimulus as more negative than the safe stimulus (C). IU was not significantly related to arousal rating difference scores between threat and safe stimuli (D). Valence, 1=negative and 9 = positive; Arousal 1 = calm, 9 = excited. Bars represent standard error.

For the valence rating difference score (Threat cue – Safe cue), STAI-T made no significant contribution to the model at the first step [*R*^2^=.044, *F*=1.808], whilst adding IU improved the hierarchal model at trend in the second step [Δ*R*^2^ =.086, *F*(1,38)=3.746, *p*=.06].

### SCR

SCR was greater to threat (*M* = .29, *SD* = .11) vs. safe (*M* = .16, *SD* = .11) trials [Condition: *F*(1,38) = 43.815, *p* < .001]. Higher IU was related to greater SCR to threat vs. safe trials, however, this relationship was not significant [Condition x IU: *F*(1,38) = 3.059, *p* = .088].

### fMRI

For all participants threat vs. safe cues induced greater activation in the bilateral amygdala, insula, frontal operculum, pre and postcentral gyrus, paracingulate, cingulate, supramarginal gyrus and middle frontal gyrus (for full list of brain regions see Table 1 & Figure 3). During threat vs. safe trial periods, activations were observed in the bilateral insula, caudate, putamen, orbital frontal cortex, supramarginal gyrus, middle frontal gyrus, thalamus, and brain stem (for full list of brain regions see Table 1 & Figure 4), The reverse contrast, safe vs. threat trial period, revealed greater activation in the bilateral hippocampus, medial cortex, superior frontal and middle frontal gyri, and precuneus (for full list of brain regions see Table 1 & Figure 4).

**Table 1.**
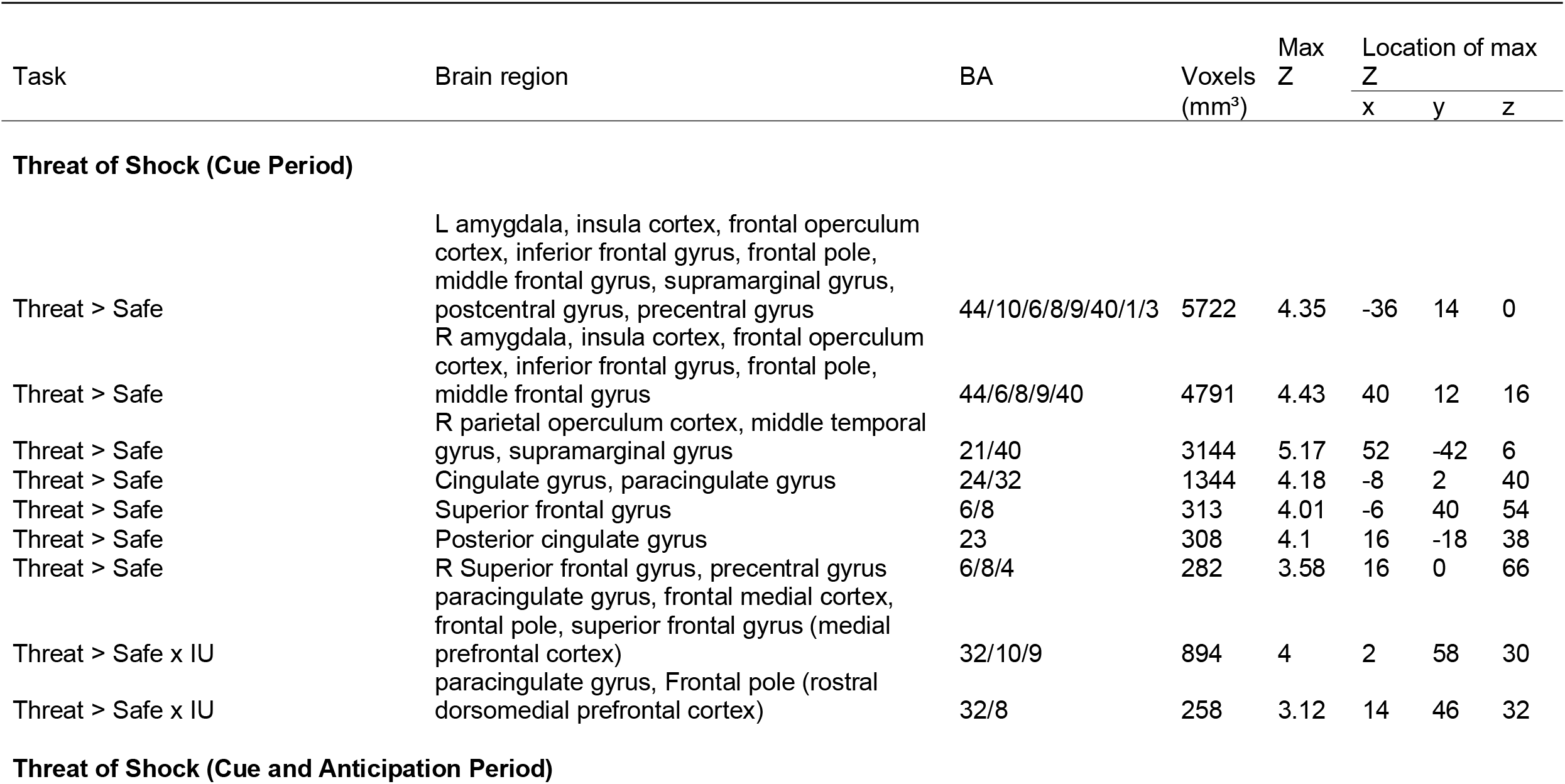
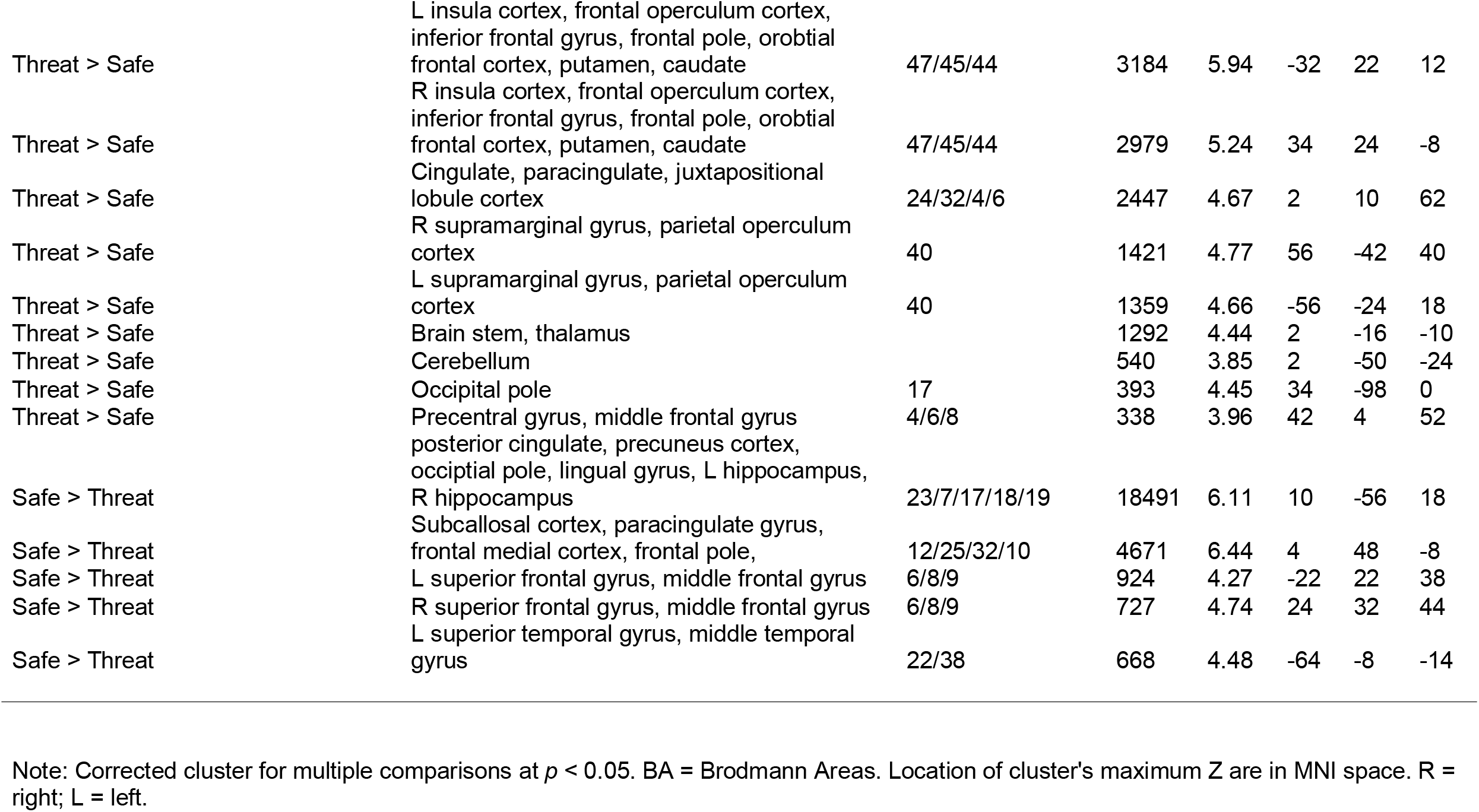
Regional activation patterns in response to stimuli presented in the threat of shock task

**Figure 3:**
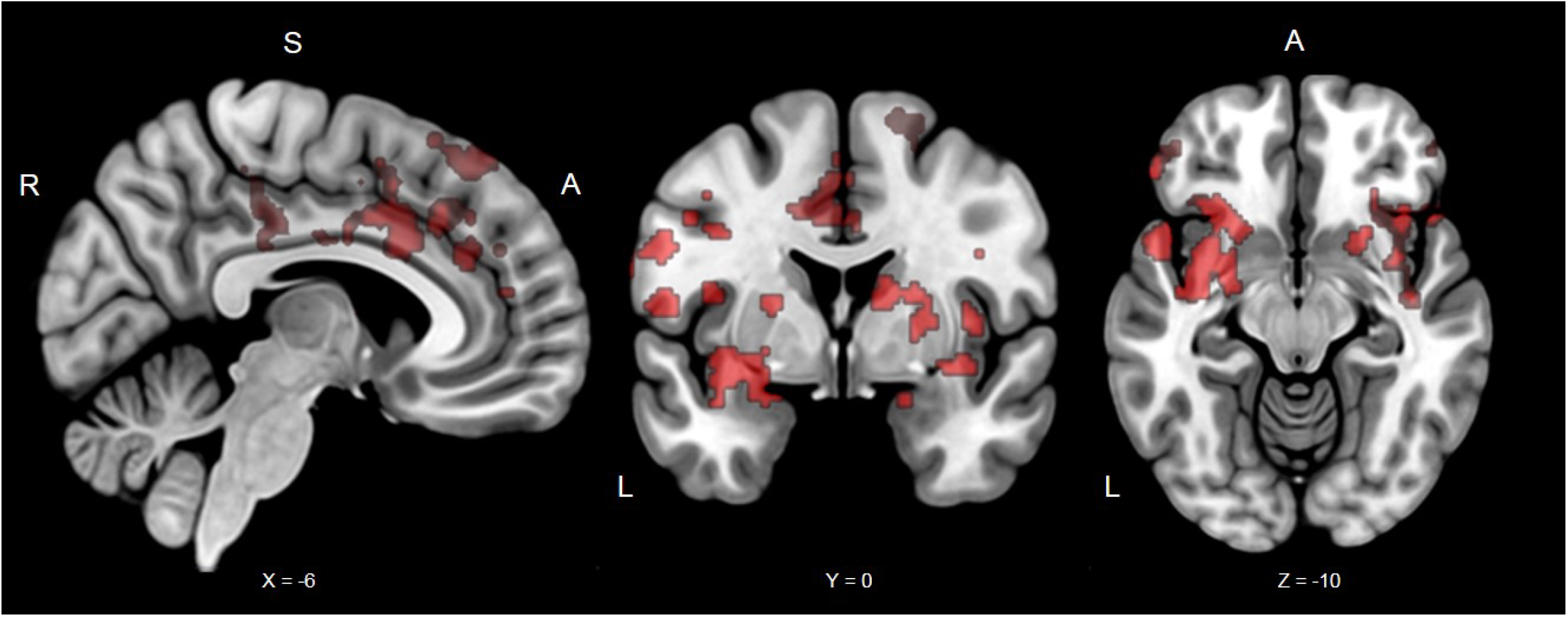
Significant clusters from the instructed threat of shock task for all participants during the cue period. Typical regions activated during threat and safety were observed. The red clusters are from the Threat > Safe contrast. Coordinates in MNI space; R, right; S, superior; A, Anterior.

**Figure 4:**
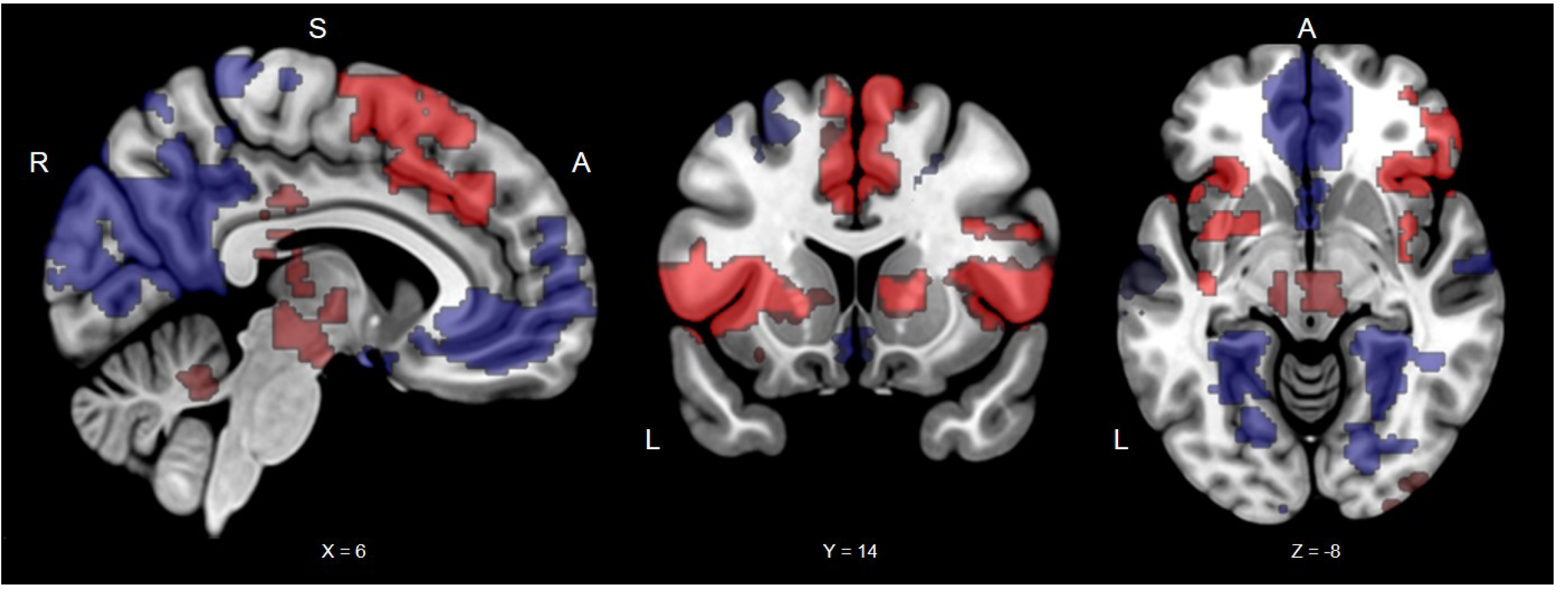
Significant clusters from the instructed threat of shock task for all participants during the cue and anticipation period. Typical regions activated during threat and safety were observed. The red clusters are from the Threat > Safe contrast and the blue clusters are from the Safe > Threat contrast. Coordinates in MNI space; R, right; S, superior; A, Anterior.

For threat vs. safe cues, high IU was associated with greater activation in the medial prefrontal cortex and rostral dorsomedial prefrontal cortex (split into two clusters, see Table 1 & Figure 5). No significant IU-related effects were observed for the safe vs. threat contrast for the cue period. In addition, no significant IU-related effects were found for the contrasts from the trial period.

**Figure 5:**
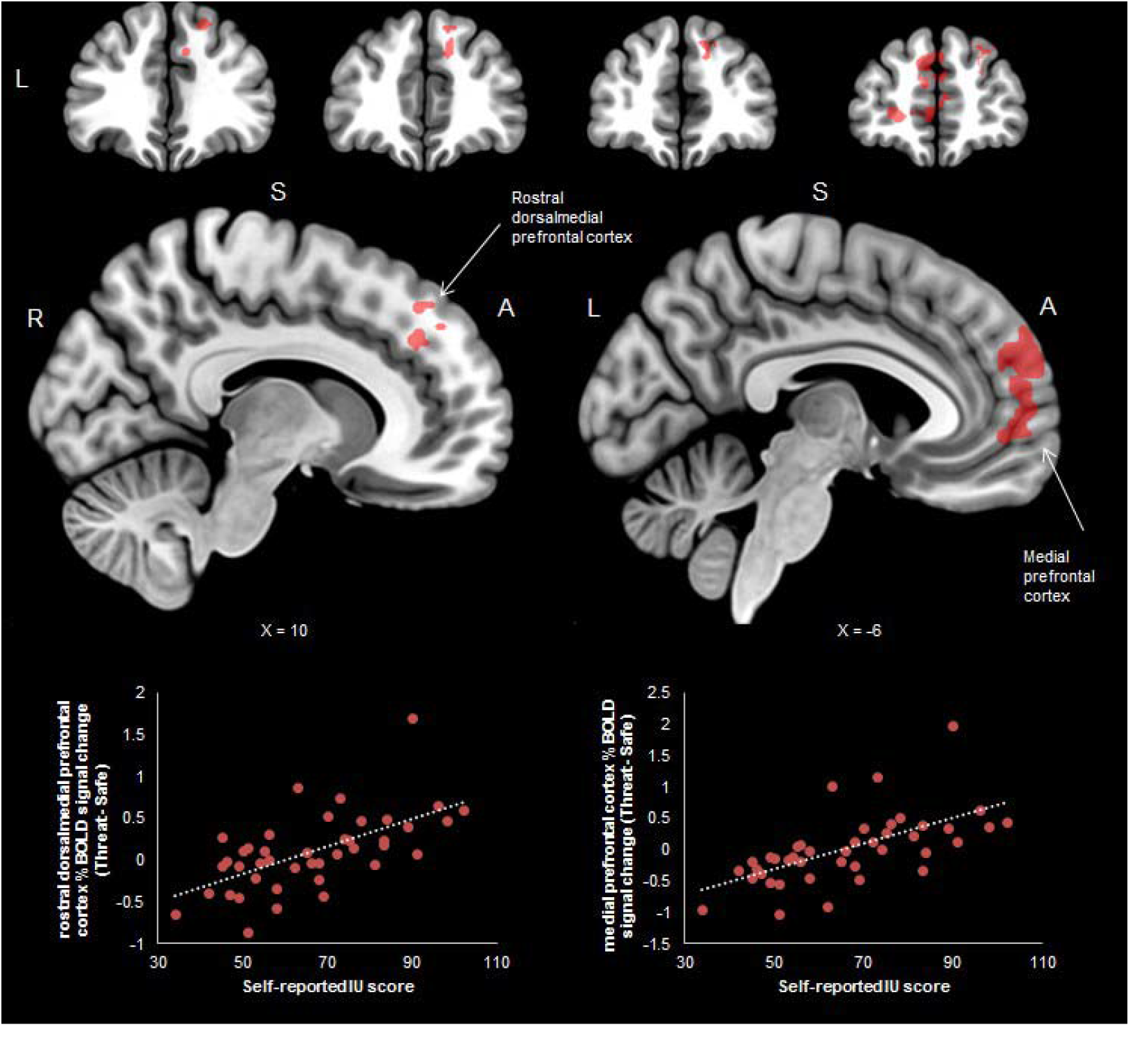
Significant clusters from the instructed threat of shock task during the cue period by individual differences in self-reported intolerance of uncertainty (IU). For threat vs. safe cues, high IU was associated with greater activation in the rostral dorsomedial prefrontal cortex and medial prefrontal cortex (see bottom of figure for correlations). Such prefrontal regions are thought to be related to safety-signalling and conscious threat appraisal. Coordinates in MNI space; R, right; S, superior; A, Anterior.

For the medial prefrontal cortex cluster (Threat cue – Safe cue), STAI-T made a significant contribution to the model at the first step [*R*^2^=.220, *F*(1,40)=11.301, *p*= .002], and adding IU in the second step improved the model significantly [Δ*R*^2^ =.167, *F*(1,39)=10.610, *p*=.002]. Similarly, for the rostral dorsomedial prefrontal cortex cluster (Threat cue – Safe cue), STAI-T made a significant contribution to the model at the first step [*R*^2^=.218, *F*(1,40)=11.171, *p*=.002], whilst adding IU improved the model significantly in the second step [Δ*R*^2^ =.163, *F*(1,39)=10.287, *p*=.003].

## Discussion

In the following experiment, self-reported IU, was not found to be associated with amygdala or insula activity during instructed outcome uncertainty of threat (i.e. if a threat will occur or not). However, higher IU was specifically associated with greater recruitment of the medial prefrontal cortex and dorsomedial rostral prefrontal cortex to cues that signalled uncertainty of threat of shock vs. safety from shock, over STAI-T. Furthermore, IU-related effects were specific to the cue (phasic); we did not observe IU modulation of neural activity during the entire trial period (sustained). These results highlight the potential of IU-based modulation of mechanisms related to safety-signalling and conscious threat appraisal in anxiety disorder pathology.

In general, we found that participants recruited typical regions associated with instructed threat of shock tasks (Etkin et al., 2011; Grupe & Nitschke, 2013; Levy & Schiller, 2021; Mechias et al., 2010; Morriss et al., 2019; Tashjian et al., 2021). Participants recruited greater amygdala to threat vs safe cues (phasic), as well as greater putamen, caudate and insula during threat vs. safe trial periods (cue + anticipation window). Moreover, participants recruited greater medial prefrontal cortex during safe vs. threat periods. As expected, greater SCR was observed to the threat vs. safe trials. Furthermore, participants rated threat cues as negative and moderately arousing, and safe cues as moderately positive and low in arousal.

We did not observe higher IU to differentially engage the amygdala or insula (across the cue or entire trial period), or display greater SCR to cues that signalled uncertainty of threat of shock vs. safety from shock. Whilst this is at odds with previous research (Morriss et al., 2015; Schienle et al., 2010; Shankman et al., 2014; Simmons et al., 2008; Somerville et al., 2013), there may be an explanation for these results. In the current study, the dominant source of uncertainty was instructed outcome of uncertainty of threat (i.e. if a threat (shock) would occur or not), whereas in previous studies there have been multiple sources of uncertainty of threat (i.e. if, when and what type of negative picture would be displayed) (Schienle et al., 2010; Shankman et al., 2014; Simmons et al., 2008; Somerville et al., 2013). Many parameters of uncertainty in combination may be perceived as more arousing and threatening in general, but particularly in individuals who score higher in IU. Therefore, in a context where different parameters of uncertainty are combined, threat cues are more likely to engage circuitry such as the amygdala and insula, and arousal-based physiology such as SCR, to alert the individual to this particular situation of threat. Our results and previous research need to be further explored and replicated, in order to fully understand how IU modulates neural circuitry under different parameters of uncertainty of threat, ideally in a single study where different parameters of uncertainty of threat are isolated (i.e., if, when and what; instructed vs. uninstructed) (Bennett, Dickmann, & Larson, 2018; Davies & Craske, 2015; Mertens & Morriss, 2021; Morriss, Bennett, & Larson, 2021; Morriss, Biagi, & Dodd, 2020).

The medial prefrontal cortex has been implicated in threat regulation and safety-signalling (Etkin et al., 2011; Milad & Quirk, 2012; Tashjian et al., 2021). In this task, higher IU was associated with greater recruitment of the medial prefrontal cortex during threat vs. safe cues. Given that the contingencies were instructed, this finding may reflect attempts to update the value of the threat cue as less threatening or safe in individuals with high IU. The modulation of activity in this area by IU is in line with prior work showing high IU individuals to recruit more medial prefrontal cortex during the extinction of threat vs. safe cues, where the values of cues change from threat to safe (Morriss et al., 2015). Higher IU was also associated with greater dorsomedial rostral prefrontal cortex to cues signalling uncertainty of threat of shock vs. safety from shock. In a recent meta-analysis, the dorsomedial rostral prefrontal cortex has been suggested to underlie conscious threat appraisal during instructed threat conditioning (Mechias et al., 2010) and generally involved in estimating threat (Grupe & Nitschke, 2013). Therefore, in the context of instructed threat of shock, greater engagement of the dorsomedial rostral prefrontal cortex may reflect conscious threat appraisal in individuals high in IU. Alongside these neural findings, we also observed individuals with high IU, relative to low IU, to rate the threat cue as more aversive. The IU-related effects for the ratings provide further evidence that individuals with higher IU found the threat cue aversive, despite being instructed about threat and safe contingencies.

Although unexpected, IU-related effects in the medial prefrontal and dorsomedial rostral prefrontal cortex were only observed for the cue period (and not the trial period) during the instructed threat of shock task. Tentatively, these findings suggest that individuals with high IU may find the cue period to be particularly salient, as this is the point at which estimates of threat and safety can be computed. Perhaps, in the context of instructed threat of shock, individuals with high IU, relative to low IU are motivated to identify the cue, in order to estimate threat and safety based on previous contingency instruction. However, once the cue is identified and the ‘risk’ is known, individuals with high IU show similar anticipation of the outcome to that of individuals with low IU.

In neurobiological models of uncertainty and anticipation (Grupe & Nitschke, 2013) the medial and dorsomedial rostral prefrontal cortex are thought to be involved in estimating threat and uncertainty, and signalling safety respectively. Notably, evidence-based therapies such as Cognitive Behavioural Therapy aim to improve flexibility in estimates of threat, safety and uncertainty in anxious populations (Clark & Beck, 2011; Saklovskis, 1996). Therefore, the IU-based modulation of neurocircuitry implicated in estimates of threat, safety and uncertainty within this study demonstrates promise for IU as a fundamental dimension in neurobiological models of uncertainty and anticipation (Grupe & Nitschke, 2013; Peters et al., 2017), as well as a potentially useful transdiagnostic treatment target in evidence-based therapies such as CBT (Dugas et al., 2003; Oglesby, Allan, & Schmidt, 2017; Robichaud & Dugas, 2006).

The study did have a few shortcomings and limitations. Firstly, while the study did include practise trials that provided participants with experience of the temporal structure of the task (i.e., anticipation time before shock), the study did not use an explicit countdown to the shock (Shankman et al., 2014; Somerville et al., 2013), which could have been beneficial to remove any additional sources of temporal uncertainty of threat. Secondly, the generality of these IU-related findings should be tested in future studies using aversive stimuli other than shocks and with different reinforcement rates of outcome uncertainty of threat (Chin, Nelson, Jackson, & Hajcak, 2016). Lastly, the sample was relatively small and contained only female participants. Future studies should aim to recruit larger samples from more diverse community or clinical samples (Hiser, Schneider, & Koenigs, 2021).

Taken together, these results suggest that, during instructed outcome uncertainty of threat, IU is specifically related over STAI-T to activation in prefrontal cortical regions. These preliminary findings highlight the potential of IU in altering safety-signalling and conscious threat appraisal mechanisms in anxiety disorder pathology (Carleton, 2016a, 2016b; Grupe & Nitschke, 2013). Further research is needed to explore the generalisability of IU-related effects during instructed outcome uncertainty of threat, and how individual differences in IU modulate different parameters of uncertainty of threat (i.e. if, when and what, as well as instructed vs. uninstructed).

## Acknowledgements

This research was supported by funding from the University of Reading and by a NARSAD Young Investigator Grant from the Brain & Behavior Research Foundation (27567) and an ESRC New Investigator Grant (ES/R01145/1) awarded to Jayne Morriss. The authors thank the participants who took part in this study and members of the CINN for their advice.

