## supplementary material for "Intolerance of uncertainty is associated with heightened responding in the prefrontal cortex during instructed outcome uncertainty of threat"

**fMRI data using permutation testing**

The default cluster forming threshold value is z = 2.3 in the FSL package for fMRI data. The use of cluster forming thresholds with parametric statistics has been argued to lead to greater chance of false positives. Non-parametric approaches have been demonstrated as robust and valid, as they are based upon actual NULL data rather than a model of such data (Eklund, Nichols and Knutsson, 2016). Therefore, to test the robustness of our results, we reran our analyses using a non-parametric permutation test in FSL (i.e. Randomise with Threshold Free Cluster Enhancement (TFCE) using 5000 samples and corrected at *p* <0.05).

The results from the original analysis were similar to the permutation test. The prefrontal clusters identified from the condition x IU interaction in the original analysis were no longer significant under the permutation test (i.e. did not survive multiple comparisons). However, the same prefrontal clusters were still observable under threshold. We believe that the results from the original analysis are genuine, given the similarity between the original analysis and the permutation test.

**Analyses across time**

We examined the effect of time in our fMRI and SCR data by splitting trials into Early (9 Threat and 9 Safe trials) and Late (9 Threat and 9 Safe trials).

For the fMRI analysis we examined the interaction between Condition by Time i.e. ([Threat – Safe_Early_] – [Threat – Safe_Late_]). Second-level GLM analysis consisted of regressors for the group mean and a linear regressor for demeaned IU using FSL's OLS procedure with a cluster thresholding of *z* = 2.3 and a corrected *p* < 0.05. We did not observe any significant clusters for the interaction of Condition x Time or Condition x Time x IU.

For the SCR data, we conducted a 2 Condition (Threat, Safe) x 2 (Early, Late) x IU ANCOVA. None of the interactions were significant [Condition x Time: *F*(1,38) = 3.083, *p* = .087; Condition x Time x IU: *F*(1,38) = 3.208, *p* = .081].

**Psychophysiological interaction analysis**

We used the medial prefrontal cortex cluster as a seed region for the instructed threat of shock task. Second-level GLM analysis consisted of regressors for the group mean and a linear regressor for demeaned IU using FSL's OLS procedure with a cluster thresholding of *z* = 2.3 and a corrected *p* < 0.05. We did not observe any significant clusters for the medial prefrontal cortex seed. The lack of connectivity results may have been due to our event-related design, which is not optimal for connectivity analysis (Cisler, Bush & Steele, 2014).
